# Triple network dynamics and future alcohol consumption in adolescents

**DOI:** 10.1101/2024.11.22.624880

**Authors:** Clayton C. McIntyre, Mohammadreza Khodaei, Robert G. Lyday, Jeffrey L. Weiner, Paul J. Laurienti, Heather M. Shappell

## Abstract

**Background:** Human neuroimaging increasingly suggests that the brain is best modeled as a highly interconnected and dynamic system. However, novel methodology for studying functional brain network dynamics have never been applied to the study of adolescent alcohol consumption. We sought to determine whether brain network dynamics are related to future drinking behavior in teenagers.

**Methods:** Resting-state functional magnetic resonance imaging (fMRI) time series from 17-year-old non/low drinking participants (n=295) of the National Consortium on Alcohol and NeuroDevelopment in Adolescence (NCANDA) study were used to fit a Hidden Semi-Markov Model (HSMM). Regions of the default mode network (DMN), salience network (SN), and central executive network (CEN), collectively known as the Triple Network, were included in modeling. The HSMM identified the most-likely state sequence for each participant, a trajectory through distinct brain network states over the course of their fMRI scan. Poisson regression models were used to assess relationships between state sequence metrics and future drinking frequency. Potential sex differences in state sequence metrics or the relationship between sequence metrics and future drinking were assessed with permutation testing and interactions in regression models.

**Results:** No sex differences in state sequence metrics were observed. However, the relationship between occupancy times and future drinking frequency differed by sex for two brain states. In the full sample, occupancy time in a state characterized by high interconnectivity between the SN and CEN was negatively associated with drinking. Occupancy time in a separate state characterized by high activation in the DMN and SN, but low activation in the CEN, was negatively associated with future drinking.

**Conclusions:** Brain network dynamics may be useful neural markers of predisposition to drinking in adolescents. Brain states which make teens vulnerable or resilient to drinking may differ between sexes.

## Introduction

Many people initiate and escalate alcohol consumption during late adolescence (Esser et al., 2017, Jones et al., 2020, Spear, 2013). This is concerning because late adolescence is known to be a critical neurodevelopmental period during which neural systems responsible for cognitive control are still developing (Somerville and Casey, 2010). Neuroimaging studies in adolescents suggest that alcohol has neurotoxic effects which may alter development of brain structure as well as functional networks which underly emotion regulation, reward processing, and cognitive control (Jacobus and Tapert, 2013, Kirse et al., 2023, Muller-Oehring et al., 2018, Zhao et al., 2021). By disrupting development of these crucial systems, alcohol exposure may leave individuals at elevated risk for developing an alcohol use disorder (AUD) as they enter adulthood. Prevention of risky alcohol use in adolescence is therefore of the utmost importance. While studies have identified environmental and psychosocial factors which influence drinking risk with increasingly complex models over the years (Christiansen et al., 1989, Patrick and Schulenberg, 2013, Green et al., 2024), the neural correlates of drinking vulnerability remain unclear. To date, studies investigating the neurobiological predictors of moderate-to-heavy drinking onset in adolescents have focused on structural measures, such as gray matter volume and cortical thickness, and task-based activation of individual brain regions (Lees et al., 2021, Squeglia et al., 2017, Norman et al., 2011).

One of the most promising approaches for studying the neural correlates of human behavior is analysis of functional brain networks. Functional brain networks include nodes which represent brain regions and edges which represent functional connections between brain regions (Bullmore and Sporns, 2009). Functional connections are commonly quantified by finding the level of correlation between blood oxygen level dependent (BOLD)(Ogawa et al., 1990) signals from distinct brain regions. Highly correlated BOLD time series between two brain regions indicates synchronized activity and suggests that regions share a functional connection. If time series from an entire functional neuroimaging scan are used for a single correlation, the outcome is a “static” functional network which models the average functional connectivity of brain regions over the course of the full scan.

Converging evidence from static functional network analyses led to development of the Triple Network model (Menon, 2011). This model proposes that behavior is governed by three key subnetworks – the Default Mode Network (DMN)(Raichle, 2015), Salience Network (SN)(Sridharan et al., 2008), and Central Executive Network (CEN)(Marek and Dosenbach, 2018). According to this model, the SN directs attention either towards the DMN, which is crucial for self-referential thought, or towards the CEN, which is responsible for goal-oriented actions and cognitive control. Triple Network architecture of young people is related to neurodevelopmental and behavioral outcomes such as impulsivity and emotional regulation (Corr et al., 2022, Jones et al., 2023, Cao et al., 2023), which are known to be closely related to drinking behaviors (Colder and Chassin, 1997, Stautz and Cooper, 2013). Further, the relationship between childhood trauma and executive dysfunction has been shown to be mediated by alterations in functional connectivity in the SN, and these same connections predict future binge drinking in adolescents (Silveira et al., 2020). Taken together, these findings suggest that functional connectivity of the Triple Network may be a useful marker of vulnerability to future drinking in adolescents.

A limitation to static network analyses is that they cannot model temporal changes in functional brain connectivity which are crucial to many neural processes (Bassett et al., 2011, Kao et al., 2020, Shine et al., 2016, Shine and Poldrack, 2018). Dynamic networks, which model second-to-second changes in functional connectivity (Bassett and Sporns, 2017), may address this limitation. Dynamic networks have been shown to outperform static networks in predicting behavioral and neuropsychiatric outcomes (Jin et al., 2017, Mokhtari et al., 2018), suggesting that dynamic properties of brain networks may be crucial to bridging the gap between measurable brain function and behavior. A challenge with dynamic network analysis is that by adding a time dimension, the number of features for each brain increases drastically, meaning that these networks can become incomprehensible. To make brain network dynamics more interpretable, methods have been developed to track the brain’s trajectory through distinct functional connectivity “states” using neuroimaging time series data (Allen et al., 2014, Shappell et al., 2019). With this state-sequence-based approach, a participant’s functional connectivity at any point during a scan is represented by one of a small number of states which are shared across a sample of participants, allowing for direct comparison of the brain network dynamics between individuals or groups. Further, participants’ brain network dynamics can be statistically related to behavioral outcomes of interest (Shappell et al., 2021), including future alcohol consumption.

In this study, resting-state functional magnetic resonance imaging (fMRI) time series data from a large sample of non/low-drinking adolescents was used to relate Triple Network dynamics to self-reported alcohol consumption from the date of their fMRI scan to their one-year follow-up visit from the scan. Given the large and rapidly growing body of evidence that sex differences are an important factor to consider in studies of alcohol-related behaviors and brain function (Hammerslag and Gulley, 2016, Geels et al., 2012, Peltier et al., 2019, Agabio et al., 2017), potential sex differences were evaluated at each step of analysis.

## Methods

### Parent Study and Sample Selection

All data was collected from the National Consortium on Alcohol and NeuroDevelopment in Adolescence (NCANDA) study (Brown et al., 2015). NCANDA is a longitudinal, multisite study of 831 participants aged 12-22 years at baseline. At each study visit, participants were interviewed about their drinking behavior and completed structural and resting-state functional MRI brain scans. The NCANDA study has previously used criteria in Table 1 (adapted from (NIAAA, 2011)) to classify participants as non/low versus risky drinkers (Brown et al., 2015). No/low versus risky drinking designations can be made at any participant’s visit given that they completed the Customary Drinking and Drug Use Record (CDDR)(Brown et al., 1998) questionnaire at each prior NCANDA visit. In the present study, the primary focus was identifying patterns in brain network dynamics from the fMRI scan that are related to drinking behavior in the year following the scan. To minimize confounding age effects, a one-year age range for analyzing participants was selected. To choose a specific one-year age range, the most common age at which participants transitioned from non/low drinking to risky drinking was identified. This age was found to be when participants were 17 at the fMRI visit and approximately 18 at the subsequent annual visit. Therefore, all brain imaging data in the present analyses are from NCANDA study visits in which the participant was 17 years old and had self-reported drinking levels that did not exceed risky drinking criteria. To be included in analyses, participants also needed to have follow-up CDDR responses no more than 1.5 years following their age 17 fMRI scan and have fMRI scan data which passed quality control. These criteria resulted in 295 participants being included in analyses.

**Table 1.** NCANDA Hazardous Drinking Criteria.

| Age (years) | Lifetime days drinking | Maximum drinks on one occasion |  |
| --- | --- | --- | --- |
|  |  | Male | Female |
| 17-17.9 | >23 | >4 | >3 |
| 18-19.9 | >51 | >4 | >3 |

NIAAA minimum criteria for being labeled a hazardous drinker based on age and sex. At the age 17 visit (when participants were scanned), all participants reported drinking levels below the hazardous drinking threshold. At their subsequent annual visit, 80 participants reported drinking levels that exceeded the *lifetime days drinking* and/or the *maximum drinks on one occasion* criteria.

### MRI Acquisition and Preprocessing

Brain image acquisition for the NCANDA study has been described in detail in previous work (Muller-Oehring et al., 2018, Zhao et al., 2021). Briefly, three study sites used a 3T GE Discovery MR750 scanner and two sites used a 3T Siemens TIM TRIO scanner. For sites with GE scanners, high resolution (0.9375 mm × 0.9375 mm × 1.2 mm) T1-weighted structural scans were acquired using an Inversion Recovery-SPoiled Gradient Recalled (IR-SPGR) echo sequence (repetition time [TR] = 5.912 ms, echo time [TE] = 1.932 ms, 146 slices, acquisition time = 7 m 14 s). For sites with Siemens scanners, high resolution (0.9375 mm × 0.9375 mm × 1.2 mm) T1-weighted structural scans were acquired using a Magnetization-Prepared RApid Gradient Echo (MPRAGE) sequence (TR = 1900 ms, TE = 900 ms, 160 slices, acquisition time = 8 m 8 s). For both scanner types, resting state BOLD-weighted image sequences were acquired using a gradient-recalled echo-planar imaging sequence (TR = 2200 ms, TE=30 ms, 32 slices, acquisition time = 10 m 03 s).

All image processing was completed in SPM 12 (https://www.fil.ion.ucl.ac.uk/spm/). Image processing began with unified segmentation (Ashburner and Friston, 2005) of T1-weighted structural images using standard six tissue priors while simultaneously warping images to Montreal Neurological Institute (MNI) standard space. The first 5 volumes (11.0 s) of the functional scan were removed to allow the signal to reach equilibrium. The remaining 269 volumes were corrected for slice-time differences, then realigned to the first volume. Functional images were warped to each participants T1 image, then subsequently warped to MNI space. To remove physiological noise and low frequency drift, functional data was filtered using a band-pass filter (0.009 – 0.08 Hz). An additional 5 volumes from the beginning and ending of the scan were removed due to artifacts on the edges of the time series from the band pass filter, leaving 259 volumes. Average whole brain gray matter, white matter, and cerebrospinal fluid mean signals, along with the six degrees of freedom motion parameters obtained from realignment were regressed from the functional data. ICA-AROMA (Pruim et al., 2015a, Pruim et al., 2015b) was used to remove motion-related noise from fMRI time series data. Following SPM processing, functional images were parcellated into the functionally defined 268-region Shen atlas (Shen et al., 2013). Analyses were focused on 104 regions of the DMN, CEN, and SN of the Triple Network. 12 of 307 scans failed to pass quality control due to issues with segmentation, warping, and/or realignment, and were excluded from analyses.

### Hidden Semi-Markov Model

An HSMM (Figure 1)(Shappell et al., 2019) was applied to the full preprocessed data set of 295 scans. Each participant’s region of interest (ROI) time series data, denoted as *Y*_*i*1_, …, *Y*_*iT*_, consisted of a *p*–dimensional vector *Y_it_* which contained the BOLD measurements of the *p* ROIs at the *t^th^* timepoint for the *i^th^* participant. The collection of vectors of observed time-series data is denoted as ^*Y*^^_*i*_. A hidden network index variable underlying the observed time-series vectors at a given observation time is denoted by *S*_*it*_, and the vector of true network state variables *S*_*i*1_, …, *S*_*iT*_ is represented by ^*S*^^_*i*_.

**Figure 1.**
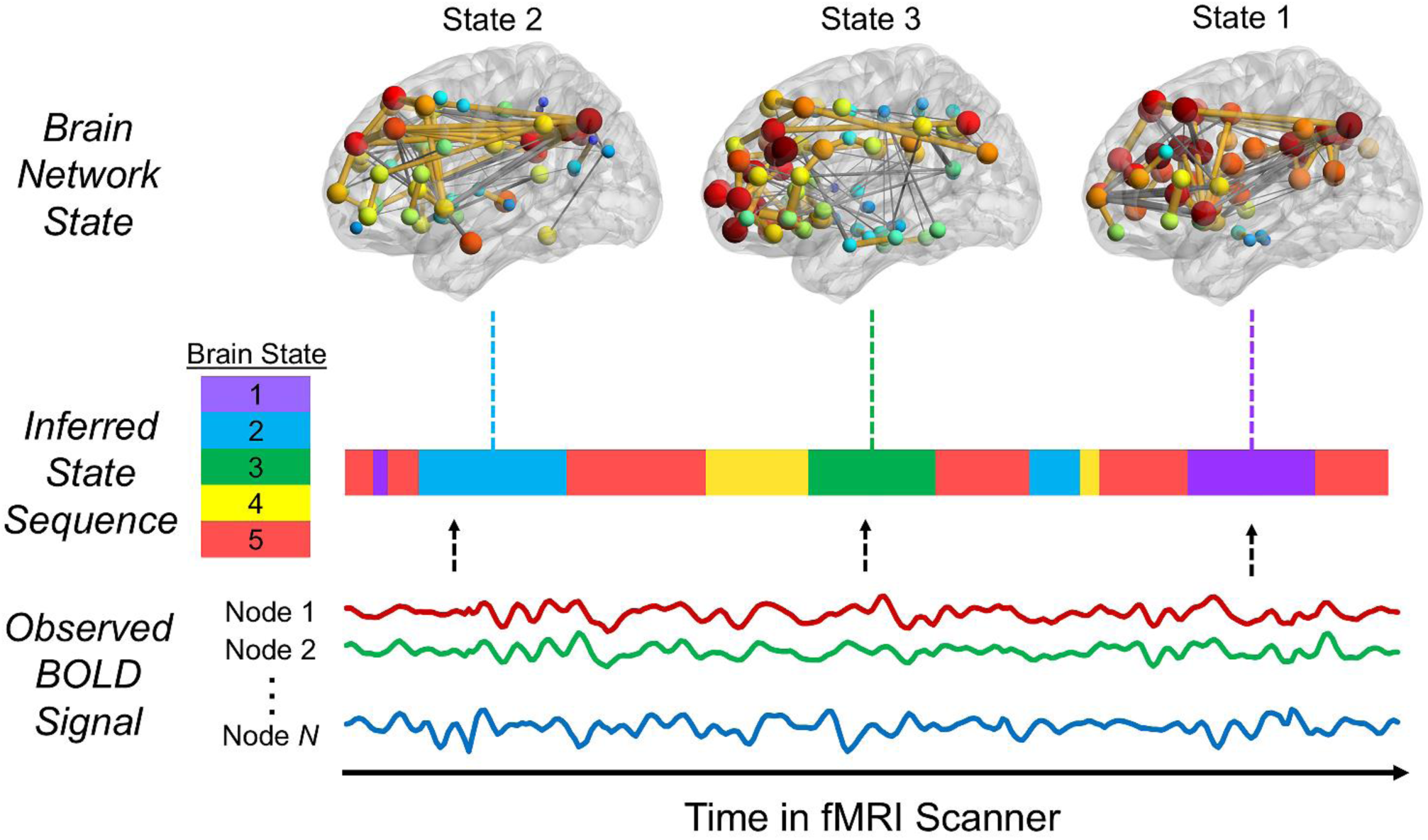
Hidden Semi-Markov Model overview. The Hidden Semi-Markov Model infers participant state sequences from observed BOLD signal time series. State sequences depict each participant’s most likely brain network state at each timepoint of their fMRI scan.

Each *Y_it_* was assumed to follow a multivariate Gaussian distribution *Yit*∼ *N*(μ_s=k_, ∑_s=k_) where the mean and covariance are dependent on the current (unknown) network state *k* of the timepoint. Thus, each network state has its own set of mean activations across our pre-specified ROIs, as well as its own covariance/correlation structure between ROIs. One set of network states was fit using the time series data across all individuals. The number of states one can fit must be specified *a priori* and is heavily dependent on the number of participants, scan timepoints, and number of network nodes included in analyses. This is because as the number of states increases, the number of parameters required to be estimated also increases. For the data included in the present analyses, five states produced stable parameter estimates over multiple fittings of the model.

The true network variables evolve over time and are modeled via a conditional transition probability matrix and a sojourn distribution. The transition probabilities model which network state a participant enters after leaving their current network state. The sojourn distribution models the probability of remaining in a given state once it is entered. A smoothed-nonparametric sojourn distribution was modeled since the exact form of the sojourn distribution was not known.

### Maximum Likelihood Estimation

HSMM parameters (each state’s mean vector and covariance matrix, initial state probabilities, sojourn distributions, and state transition probabilities) were estimated utilizing the *mhsmm* R package (O’Connell and Hojsgaard, 2011). This package allows for inferring multiple observation sequences (i.e., multiple participants at once). Therefore, each individual’s time-series data was concatenated so that the total number of rows in our data set was equal to the sum of the time-series lengths across all individuals. The number of columns in our data set was equal to the number of regions included (*p* = 104). The HSMM parameters were then estimated on this entire data set, resulting in one set of model parameter estimates.

The Expectation-Maximization algorithm (Dempster et al., 1977) was used to estimate all model parameters. Each participant’s most probable sequence of true network states was estimated using the Viterbi Algorithm (Forney, 1973). The Viterbi algorithm takes state estimates, transition probability estimates, sojourn density estimates, and initial state probability estimates obtained from fitting the HSMM on all participants and calculates each individual participant’s most likely sequence of states given those estimates and the participant’s BOLD time-series data. Said differently, the Viterbi algorithm takes an individual’s time-series data and determines which of five states (estimated from the entire data set) the participant is in at each timepoint during their scan. Therefore, although state estimates are based on the entire data set and are not subject-specific, any given subject’s state sequence is specific to them. A further description of the model estimation procedure, including how the non-parametric sojourn densities were estimated, can be found in previous work (Shappell et al., 2019).

### Network Community Assignment Inference

To characterize the network topology of the five functional connectivity states identified by the HSMM, covariance matrices (estimated via the maximum likelihood estimation routine described above) were converted to correlation matrices, then used as an undirected, weighted adjacency matrix. The Louvain community detection algorithm from the Brain Connectivity Toolbox (www.brain-connectivity-toolbox.net)(Rubinov and Sporns, 2010) was used to identify modules for each state. A module is a grouping of nodes which are more connected to each other than they are to nodes of other modules (Blondel et al., 2008, Newman and Girvan, 2004). For each state, the Louvain algorithm was run for 1,000 iterations. The module assignments of the iteration with the highest *Q* (a scalar quantification of a network’s modularity (Newman and Girvan, 2004)) was designated as the state’s community structure.

### Group Difference Permutation Testing

We assessed whether there were group differences in empirical sojourn distributions of those who went on to exceed risky drinking criteria for non/low drinking at their subsequent visit (n = 80) and those who did not (n = 215). Additionally, HSMM metrics of males (n = 143) and females (n = 152) were compared. To find empirical sojourn distributions for each state, the number of consecutive time points spent in states for each individual sojourn was counted in each participant. These counts were then compiled across all participants for each group. These group-level counts were then converted into a density estimate for each group. Each participant’s initial state and last state were omitted from these counts as it is impossible to know how long they were in the initial state prior to the beginning of their fMRI scan or how long they remained in the final state following the conclusion of fMRI data collection. Therefore, omitting these counts prevented biasing the sojourn distributions towards shorter dwell times.

Once separate empirical sojourn distributions were obtained for each group for all five states, a group comparison of each state’s empirical sojourn distribution was performed using Kullback-Leibler (KL) divergence, a measure of the directed divergence between two probability distributions (Csiszar, 1975, Kullback and Leibler, 1951). For a more detailed discussion of this approach, see (Shappell et al., 2019). The *Philentropy* R package was used to calculate KL divergence. To determine if there were any group differences in sojourn distributions in each state, permutation testing with 1,000 permuted samples were used.

### Poisson Outcomes and Relationships with Future Alcohol Consumption

In addition to group comparisons, relationships between HSMM model outputs and responses to the CDDR “Since your last visit, how many days did you drink alcohol?” were assessed for statistical significance. This question was chosen because it assesses the regularity of drinking behavior and mirrors the frequency aspect of the NIAAA’s risky drinking criteria. Responses to this question were skewed towards fewer drinking occasions (i.e., most participants reported drinking very rarely). In rare cases where participants responded with non-integer values to the number of days they drank alcohol, responses were rounded to the nearest whole number.

We found that future drinking outcomes were zero-inflated due to many participants (148 out of 295) reporting no drinking in the year between visits. To address this, two-stage regression analyses were used to identify relationships between state sequence metrics and future drinking days. The first stage was a binomial regression that determined whether there were statistical differences in occupancy time or the number of sojourns into each state for participants who reported no drinking versus participants who reported drinking at least once. The second stage was a Poisson regression model which included only participants who reported drinking at least once between visits. The Poisson regression model determined whether occupancy time and/or number of sojourns into each state was significantly related to the number of days drinking between visits.

To determine whether any findings were sex-dependent, we tested for significant interactions between sex and HSMM metrics in regression models with CDDR responses as outcomes. Ten total models were fit (two for each state: one with occupancy time as the independent variable and one with number of sojourns as the independent variable). Number of drinking days was the outcome of interest. Because many separate models were fit, all *p*-values were adjusted for multiple comparisons using the Bonferroni approach (Bland and Altman, 1995, Dunn, 1961). In cases where significant sex interactions were identified, subsequent sex-stratified regression models were fit to assess sex-dependent relationships between HSMM metrics and drinking outcomes. In models where no significant sex interaction was identified, the main effect across males and females together is reported. All models were adjusted for drinking levels prior to the fMRI scan, time between study visits, household income, study site, and motion during the fMRI scan (both maximum spike and mean framewise displacement). Models with number of sojourns as the independent variable of interest additionally controlled for the total number of sojourns across all states. For all analyses, findings were considered statistically significant at a familywise error rate of 0.05.

## Results

### Sample Characteristics and Drinking Levels

Of the 295 participants included in network analyses, 152 (51.6%) were female, 216 (73.2%) identified as white, and 33 (11.2%) identified as Hispanic. At the age 17 visit, the average age was 17.5 ± 0.3. At the subsequent annual visit, the average age was 18.5 ± 0.3. Figure 2 shows each participant’s reported number of days drinking in their lifetime prior to their age 17 visit and between study visits. Among people who consumed any alcohol before the age 17 visit, the mean number of lifetime days drinking before the visit was 4.3 ± 3.7. Among people who consumed any alcohol between visits, the mean number of days drinking between visits was 10.3 ± 14.6.

**Figure 2.**
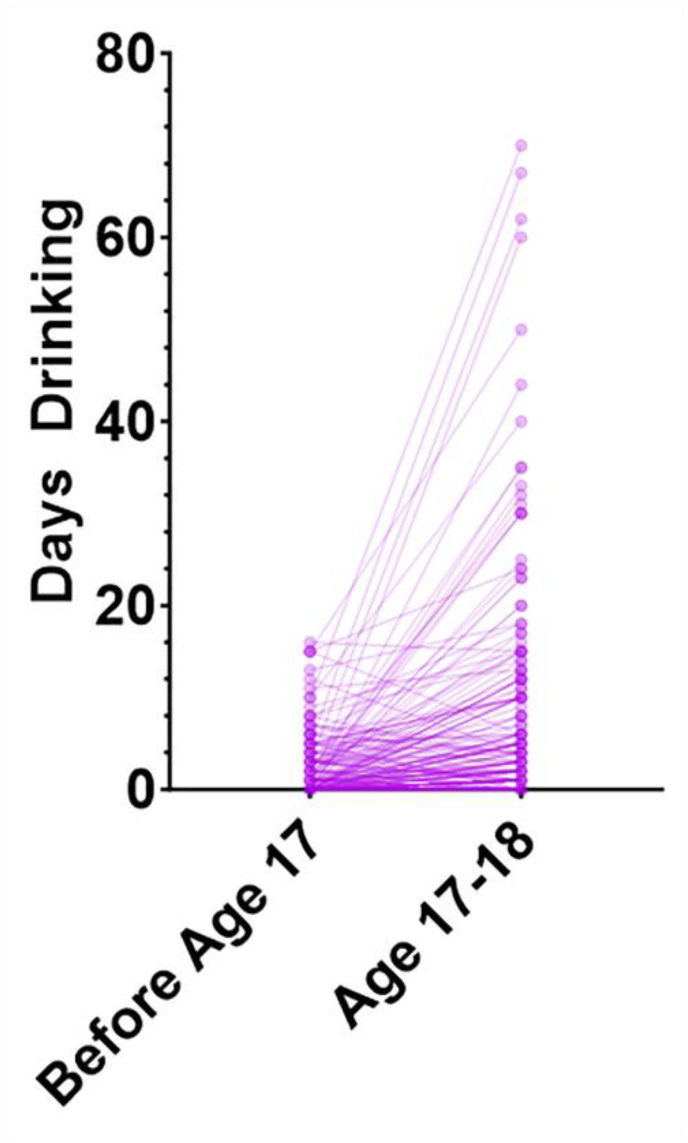
Drinking days before and after fMRI scan Each dot represents an individual participant’s visit. Lines connect questionnaire responses from two different visits. On the left (“Before Age 17”), the sum of lifetime drinking days prior to the age 17 NCANDA visit is shown. On the right (“Age 17-18”), the number of days drinking between the two NCANDA visits is shown. 80% (124 of 155) of participants who drank any alcohol consumption by age 18 reported that between the age 17 and 18 visits, they consumed alcohol on as many or more days as they had in their lifetime before age 17.

Two sample t-tests were used to compare drinking frequency in males and females who reported any drinking. Among males (n = 36) and females (n = 49) who had reported any drinking in their lifetime prior to the age 17 visit, there was no difference in the number of days alcohol was consumed (*p* = 0.996). Among males (n = 72) and females (n = 75) who reported any drinking in the year between visits, there was again no difference in the number of days alcohol was consumed (*p* = 0.662).

### Functional Brain State Characterization

Correlation matrices of each brain state are shown in Supplementary Figure 1. Figure 3 shows *Q* values for each state. States 2 and 3 had the highest modularity whereas State 4 had the lowest modularity. Node community assignments for each state are mapped into brain space in Figure 4 along with a reference image of expected community assignments based on Triple Network connectivity patterns. The red, yellow, and blue node coloration corresponds roughly to the DMN, SN, and CEN, respectively. While nodes in States 2-5 are very consistent with the Triple Network, State 1 deviates from this pattern in that regions which are typically members of the SN (i.e., the anterior insula, anterior/mid cingulate, and amygdala) appear in the same network module (i.e., the blue module) as regions of the CEN in this state. The yellow module in State 1 is mostly limited to the thalamus and basal ganglia.

**Figure 3.**
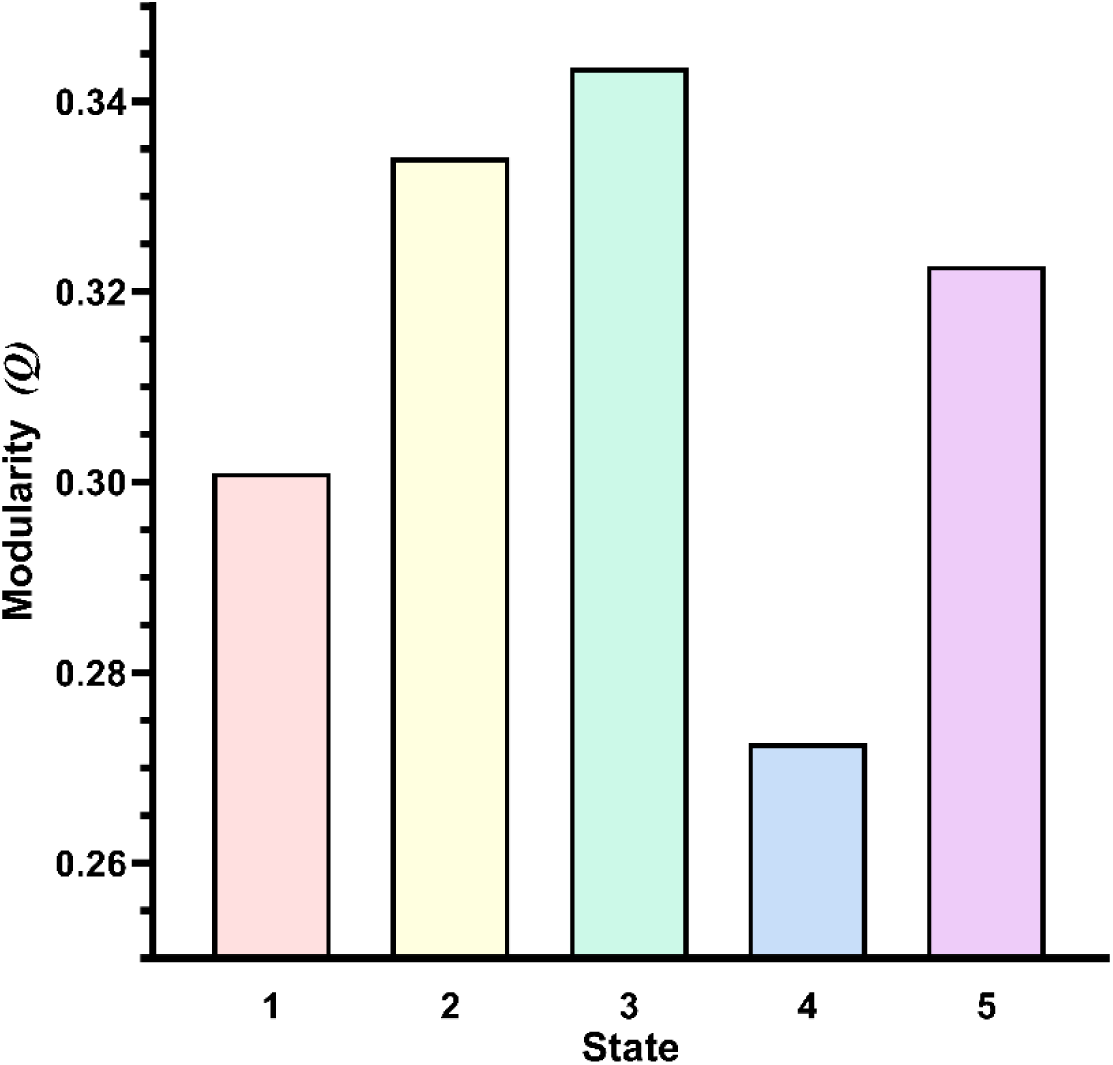
State modularity level Each brain state’s modularity level is quantified using the scalar *Q*. States 2 and 3 had the highest modularity (i.e., more connections within communities and fewer connections between communities) while state 4 had the lowest modularity.

**Figure 4.**
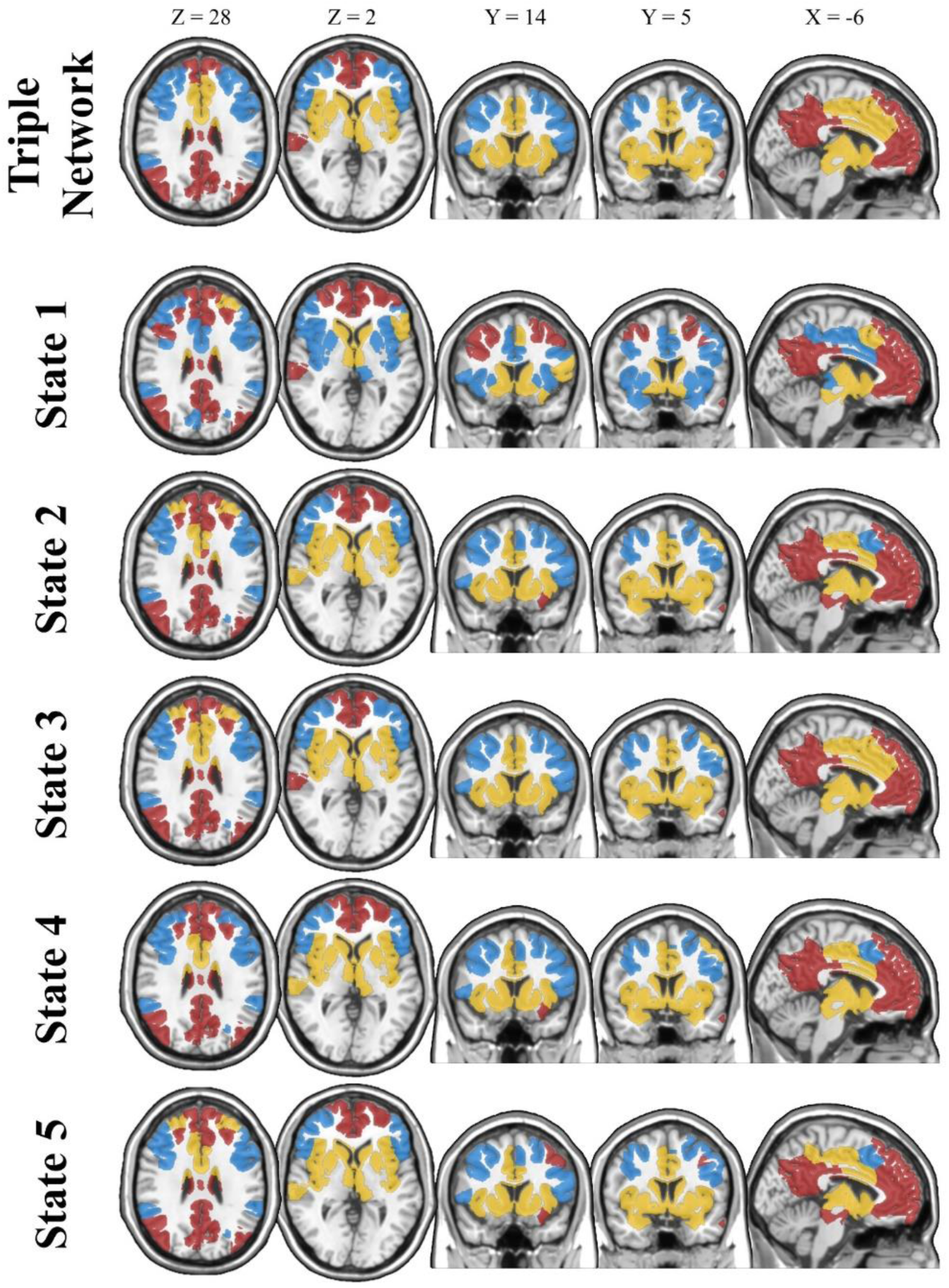
Functional network state community structure 2-dimensional brain slices depicting node community assignments in each network state. Regions with the same coloring appeared in the same network community. Each state had three network communities (shown as red, blue, and yellow). While states 2-5 followed modularity patterns that closely follow the Triple Network, state 1 deviated from this pattern in that regions which are members of the salience network (i.e., the amygdala, insula, cingulate) appeared in the same community as regions typically associated with the central executive network. MNI coordinates of brain slices are shown above each column.

Figure 5 shows the average BOLD signal of nodes in each state relative to each node’s average BOLD signal over the full scan. State 2 demonstrated relatively high activation in regions of the CEN but low activity in the DMN. State 5 demonstrated relatively high activation in regions of the DMN and SN but low activation in the CEN. States 1, 3, and 4 all showed relatively homogenous activation across each of the Triple Network subnetworks.

**Figure 5.**
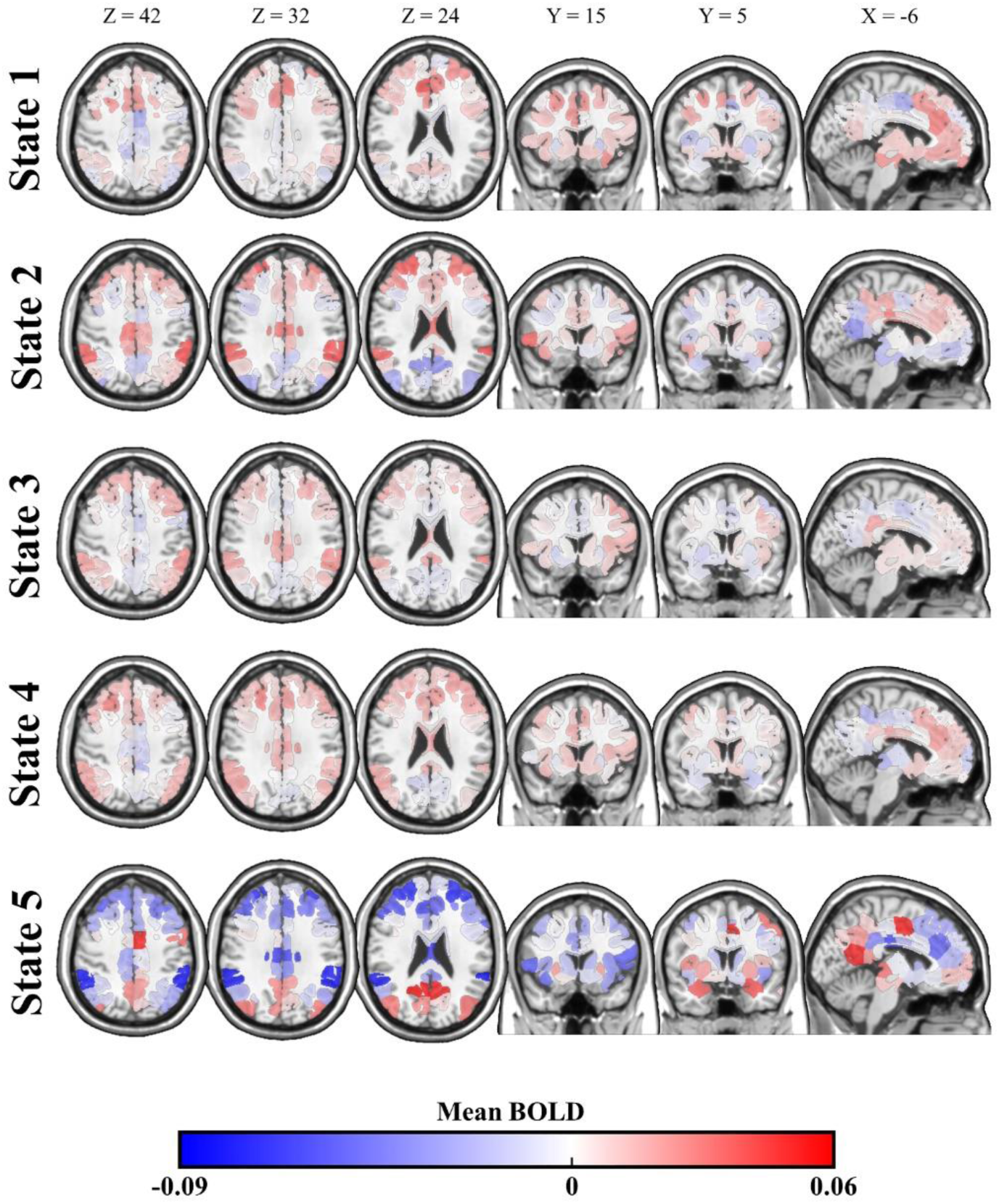
Mean BOLD of nodes in each state 2-dimensional brain slices depicting the mean activation (i.e., BOLD signal) in each network state. Activation values indicate each node’s deviation (on average in a given state) from its overall average BOLD signal from the full fMRI time series. Notably, state 2 demonstrates relatively high activation in regions of the central executive network and low activation in regions of the default mode network. State 5 demonstrates high activity in parts of the default mode and salience network with low activation in the central executive network. States 1, 3, and 4 have largely homogenous activation levels across each of the Triple Network subnetworks. MNI coordinates of brain slices are shown above each column.

### Group Comparisons of Sojourn Distributions

We compared sojourn distributions for each state between participants who did (n = 80) and did not (n = 215) exceed risky drinking threshold at their follow up visit. Permutation testing of group sojourn distributions revealed no significant differences in sojourn distributions in any states (Supplementary Table 1). We also compared sojourn distributions for each state between males (n = 143) and females (n = 152). Males and females had a slight difference in sojourn distributions for State 2 such that males tended to dwell in State 2 for longer than females (Supplementary Figure 3). However, this difference was not statistically significant following Bonferroni correction for multiple comparisons. Males and females did not differ in sojourn distribution for any other state (Supplementary Table 2).

### Relationship between HSMM Metrics and Future Drinking Days

No results from the first stage (binomial regression) of regression analyses for sex interactions and main effects were statistically significant and are shown in Supplementary Tables 3 and 4. Results from second stage analyses (Poisson regression) for sex interactions and main effects are shown in Tables 2 and 3, respectively.

**Table 2.** Stage 2 Poisson Regression Results – Sex Interactions.

| Sequence Metric | Risk Ratio | Standard Error | *p*<sub>Bonferroni</sub> |
| --- | --- | --- | --- |
| Occupancy – State 1 | 0.9988 | 0.0012 | 1 |
| Occupancy – State 2 | 0.9953 | 0.0008 | <0.001 |
| Occupancy – State 3 | 1.0001 | 0.0009 | 1 |
| Occupancy – State 4 | 1.0064 | 0.0010 | <0.001 |
| Occupancy – State 5 | 0.9978 | 0.0014 | 1 |
| Number of Sojourns – State 1 | 1.0327 | 0.0221 | 1 |
| Number of Sojourns – State 2 | 1 | 0.0142 | 1 |
| Number of Sojourns – State 3 | 1.0166 | 0.0155 | 1 |
| Number of Sojourns – State 4 | 0.9181 | 0.0204 | 0.010 |
| Number of Sojourns – State 5 | -0.9695 | 0.0141 | 1 |

Each row shows results from a different Poisson regression model, all with number of days drinking between visits as the outcome variable. Standard errors are based on the $\beta$ -values from Poisson regressions, which indicate logarithmic relationships between the independent and dependent variable. For interpretability, $\beta$ -values are reported as Risk Ratios according to the formula Risk Ratio = $e^{\beta}$ .

**Table 3.** Stage 2 Poisson Regression Results – Main and Stratified Effects.

| Sequence Metric | Risk Ratio | Standard Error | *p*<sub>Bonferroni</sub> |
| --- | --- | --- | --- |
| Occupancy – State 1 | 0.9978 | 0.0006 | **0.030** |
| Occupancy – State 2 (Male) | 1.0047 | 0.0008 | **<0.001** |
| Occupancy – State 2 (Female) | 0.9977 | 0.0007 | **0.019** |
| Occupancy – State 3 | 1.0015 | 0.0005 | **0.044** |
| Occupancy – State 4 (Male) | 0.9973 | 0.0009 | 0.073 |
| Occupancy – State 4 (Female) | 1.0038 | 0.0005 | **<0.001** |
| Occupancy – State 5 | 0.9972 | 0.0007 | **0.001** |
| Number of Sojourns – State 1 | 0.9627 | 0.0126 | 0.109 |
| Number of Sojourns – State 2 | 1.0109 | 0.0114 | 1 |
| Number of Sojourns – State 3 | 1.0002 | 0.0086 | 1 |
| Number of Sojourns – State 4 (Male) | 0.9778 | 0.0168 | 1 |
| Number of Sojourns – State 4 (Female) | 1.0934 | 0.0142 | **<0.001** |
| Number of Sojourns – State 5 | 0.9827 | 0.0115 | 1 |

In second stage analyses, significant sex interactions were identified for occupancy time in States 2 (*p_Bonferroni_ <* 0.001) and 4 (*p_Bonferroni_ <* 0.001). Additionally, a significant interaction was identified between sex and the number of sojourns into State 4 (*p_Bonferroni_* = 0.010). Sex-stratified models revealed that occupancy time in State 2 was positively associated with future drinking days in males (Risk Ratio [RR] = 1.005, SE = 0.0008, *p_Bonferroni_* < 0.001) but negatively associated with future drinking days in females (RR = 0.998, SE = 0.0007, *p_Bonferroni_* = 0.019). This means that among males, each additional TR spent in State 2 would increase the expected number of future drinking days by 0.5% if all other variables are held constant. In females, each additional TR spent in State 2 would be expected to correspond to a 0.2% decrease in drinking days. Occupancy time in State 4 was not significantly associated with future drinking days in males (RR = 0.997, SE = 0.0009, *p_Bonferroni_* = 0.073) but was positively associated with future drinking days in females (RR = 1.004, SE = 0.0005, *p_Bonferroni_* < 0.001). Additionally, the number of sojourns into state 4 was not significantly associated with future drinking days in males (RR = 0.978, SE = 0.0168, *p_Bonferroni_* = 1) but was positively associated with future drinking days in females (RR = 1.093, SE = 0.0142, *p_Bonferroni_* < 0.001).

Across the whole sample (i.e., in cases when there were no significant sex differences), the number of days drinking between visits was negatively associated with occupancy time in States 1 (RR = 0.998, SE = 0.0006, *p_Bonferroni_* = 0.030) and 5 (RR = 0.997, SE = 0.0007, *p_Bonferroni_* = 0.001). The number of drinking days between visits was positively associated with occupancy time in State 3 (RR = 1.002, SE = 0.0005, *p_Bonferroni_* = 0.044).

## Discussion

This study aimed to determine whether dynamic connectivity of the Triple Network in non/low-drinking adolescents was related to future drinking behavior. We also assessed whether biological sex influenced relationships between brain network dynamics and future drinking. Sojourn distributions for the 5 functional connectivity states that were identified did not distinguish between future risky drinkers and future non/low drinkers, but there were several significant associations between occupancy times in each state and future drinking days, either in one biological sex or across the whole sample. We discuss the implications of these findings below.

Occupancy time in State 1 was negatively associated with future drinking days across the full sample. State 1 was distinct from other brain states in that it followed a unique modularity pattern which suggested strong functional connectivity between hubs of the SN (including the anterior insula, amygdala, and anterior/mid cingulate) and the CEN. Prior work suggests that stronger connectivity between frontal regions of the CEN and striatal portions of the SN is associated with goal-oriented control and reduced compulsivity in adolescents (Vaghi et al., 2020, Vink et al., 2014). Functional connections between regions of the SN and CEN are also known to strengthen over the course of normal adolescent neurodevelopment (Somerville and Casey, 2010). This is thought to support maturation of cognitive control abilities which reduce susceptibility to engaging in risky behavior, such as underage drinking. Therefore, our finding that time spent in a state with strong SN-CEN integration is protective against future drinking is not surprising and may indicate that individuals who spend large amounts of time in this state are less vulnerable to drinking or other risky behaviors which are commonly observed in adolescents.

Menon’s original paper introducing the Triple Network model (Menon, 2011) suggested that an inability to appropriately activate or deactivate any of the three Triple Network subnetworks is likely to be associated with deleterious psychiatric/behavioral outcomes. In the absence of salient external stimuli (i.e., in resting-state scans), participants are expected to show increased activation in the DMN and SN and reduced activation of the CEN. This pattern was best represented by State 5. Occupancy time in state 5 was negatively associated with future drinking days, suggesting that time spent in this state may be protective against future drinking. State 3, which had connectivity that was very similar to State 5, demonstrated relatively homogenous activation across each of the Triple Networks. In contrast to State 5, occupancy time in State 3 was positively associated with future drinking days. This suggests that transient, aberrant activation of subnetworks may be a marker of heightened vulnerability to future drinking.

State 2 showed relatively high activation of the CEN and low activation of the DMN. Time spent in State 2 was positively associated with future drinking days in males, but negatively associated with future drinking days in females. In addition to sex differences in the relationship between State 2 and future drinking frequency, occupancy time in State 4 was positively associated with future drinking days for females but had a near-significant negative relationship with future drinking days for males. These findings add to a large body of work showing that males and females differ in neural responses to alcohol cues (Seo et al., 2011, Petit et al., 2013, Kaag et al., 2019). Recent findings suggest that this difference is no longer present once individuals have developed more severe AUD (Gerhardt et al., 2022), which may indicate that sex differences are particularly important to consider in young people who are just beginning to use alcohol. Prior work suggests that the social pressures which drive problematic alcohol consumption differs in adolescent boys and girls (Schulte et al., 2009). Whereas adolescent boys are more likely to begin drinking out of curiosity or sensation-seeking, early drinking experiences in young girls are more likely to have coping motives (Kuntsche and Muller, 2012). This study is the first to show that aspects of brain network dynamics which impact resilience and vulnerability to alcohol use differ between males and females.

The present work has some limitations. First, drinking measures in this work were from self-report data. It is possible that some participants misrepresented their drinking levels either intentionally (due to stigma surrounding underage drinking, for example) or through misremembering actual drinking levels. Second, a challenge with state-sequence-based analysis of dynamic functional networks is that there is no definite answer to how many underlying brain states exist. In the present study, five states were used for analyses as several fittings of the HSMM with different numbers of states showed that models with five states seemed to produce the most stable state sequences across the sample. However, it is plausible that the brain has more than five underlying states, even during resting scans. Third, rather than analyze whole-brain connectivity dynamics, this study focused on regions of the Triple Network because HSMMs are limited in the number of brain regions that can be included in analyses. By focusing on regions of the Triple Network, network dynamics of other brain regions were not considered in analyses for this study. Finally, this work assessed the relationship between network dynamics of resting-state fMRI data and future alcohol consumption. This work cannot answer how resting-state brain network dynamics relate to neural activity in the moments leading up to alcohol consumption. It is plausible that network dynamics in environments in which alcohol is typically consumed by adolescents differ from network dynamics in resting-state. Nevertheless, resting-state connectivity has repeatedly been shown to be a useful indicator of future behavior (Mokhtari et al., 2018, Morris et al., 2022, Stevens and Spreng, 2014, Studer et al., 2013).

This work is the first to apply novel methods for analyzing functional brain network dynamics to the study of alcohol consumption risk in adolescents. There is currently no existing literature on functional brain network dynamics and adolescent drinking to which findings from this work can be compared. It is our hope that this work will serve as a foundation for future efforts to determine how brain network dynamics contribute to adolescents’ propensity to engage in risky drinking. Future analyses should explore brain regions beyond just the Triple Network. Additionally, studies should investigate different adolescent populations, such as people who already meet risky drinking criteria at the time of their brain scans or people who co-use alcohol and other substances. Finally, an important next step will be to study longitudinal development in non-drinking versus risky drinking adolescents to determine how alcohol consumption during this critical neurodevelopmental period impacts maturation of brain network dynamics.

## Acknowledgements

All data presented in this work was collected from by the NCANDA study. NCANDA is supported by NIH grants AA021697, AA021697-04S1, AA021695, AA021692, AA021696, AA021681, AA021690, and AA021691. The NCANDA dataset is publicly available upon request (see http://www.ncanda.org/index.php). Analyses were supported by NIH grants P50AA026117 and K25EB032903.

## References

Agabio, R., Pisanu, C., Gessa, G. L. & Franconi, F. 2017. Sex Differences in Alcohol Use Disorder. Curr Med Chem, 24, 2661–2670.

Allen, E. A., Damaraju, E., Plis, S. M., Erhardt, E. B., Eichele, T. & Calhoun, V. D. 2014. Tracking whole-brain connectivity dynamics in the resting state. Cereb Cortex, 24, 663–76.

Ashburner, J. & Friston, K. J. 2005. Unified segmentation. Neuroimage, 26, 839–51.

Bassett, D. S. & Sporns, O. 2017. Network neuroscience. Nature Neuroscience, 20, 353–364.

Bassett, D. S., Wymbs, N. F., Porter, M. A., Mucha, P. J., Carlson, J. M. & Grafton, S. T. 2011. Dynamic reconfiguration of human brain networks during learning. Proc Natl Acad Sci U S A, 108, 7641–6.

Bland, J. M. & Altman, D. G. 1995. Multiple Significance Tests - the Bonferroni Method.10. British Medical Journal, 310, 170–170.

Blondel, V. D., Guillaume, J. L., Lambiotte, R. & Lefebvre, E. 2008. Fast unfolding of communities in large networks. Journal of Statistical Mechanics-Theory and Experiment.

Brown, S. A., Brumback, T., Tomlinson, K., Cummins, K., Thompson, W. K., Nagel, B. J., De Bellis, M. D., Hooper, S. R., Clark, D. B., Chung, T., Hasler, B. P., Colrain, I. M., Baker, F. C., Prouty, D., Pfefferbaum, A., Sullivan, E. V., Pohl, K. M., Rohlfing, T., Nichols, B. N., Chu, W. & Tapert, S. F. 2015. The National Consortium on Alcohol and NeuroDevelopment in Adolescence (NCANDA): A Multisite Study of Adolescent Development and Substance Use. J Stud Alcohol Drugs, 76, 895–908.

Brown, S. A., Myers, M. G., Lippke, L., Tapert, S. F., Stewart, D. G. & Vik, P. W. 1998. Psychometric evaluation of the Customary Drinking and Drug Use Record (CDDR): a measure of adolescent alcohol and drug involvement. J Stud Alcohol, 59, 427–38.

Bullmore, E. & Sporns, O. 2009. Complex brain networks: graph theoretical analysis of structural and functional systems. Nat Rev Neurosci, 10, 186–98.

Cao, W., Liu, Y., Zhong, M., Liao, H., Cai, S., Chu, J., Zheng, S., Tan, C. & Yi, J. 2023. Altered intrinsic functional network connectivity is associated with impulsivity and emotion dysregulation in drug-naive young patients with borderline personality disorder. Borderline Personal Disord Emot Dysregul, 10, 21.

Christiansen, B. A., Roehling, P. V., Smith, G. T. & Goldman, M. S. 1989. Using Alcohol Expectancies to Predict Adolescent Drinking Behavior after One Year. Journal of Consulting and Clinical Psychology, 57, 93–99.

Colder, C. R. & Chassin, L. 1997. Affectivity and impulsivity: Temperament risk for adolescent alcohol involvement. Psychology of Addictive Behaviors, 11, 83–97.

Corr, R., Glier, S., Bizzell, J., Pelletier-Baldelli, A., Campbell, A., Killian-Farrell, C. & Belger, A. 2022. Triple Network Functional Connectivity During Acute Stress in Adolescents and the Influence of Polyvictimization. Biol Psychiatry Cogn Neurosci Neuroimaging, 7, 867–875.

Csiszar, I. 1975. I-Divergence Geometry of Probability Distributions and Minimization Problems. Annals of Probability, 3, 146–158.

Dempster, A. P., Laird, N. M. & Rubin, D. B. 1977. Maximum Likelihood from Incomplete Data Via Em Algorithm. Journal of the Royal Statistical Society Series B-Methodological, 39, 1–38.

Dunn, O. J. 1961. Multiple Comparisons among Means. Journal of the American Statistical Association, 56, 52-&.

Esser, M. B., Clayton, H., Demissie, Z., Kanny, D. & Brewer, R. D. 2017. Current and Binge Drinking Among High School Students - United States, 1991-2015. MMWR Morb Mortal Wkly Rep, 66, 474–478.

Forney, G. D. 1973. Viterbi Algorithm. Proceedings of the Ieee, 61, 268–278.

Geels, L. M., Bartels, M., Van Beijsterveldt, T. C., Willemsen, G., Van Der Aa, N., Boomsma, D. I. & Vink, J. M. 2012. Trends in adolescent alcohol use: effects of age, sex and cohort on prevalence and heritability. Addiction, 107, 518–27.

Gerhardt, S., Hoffmann, S., Tan, H., Gerchen, M. F., Kirsch, P., Vollstadt-Klein, S., Kiefer, F., Bach, P. & Lenz, B. 2022. Neural cue reactivity is not stronger in male than in female patients with alcohol use disorder. Front Behav Neurosci, 16, 1039917.

Green, R., Wolf, B. J., Chen, A., Kirkland, A. E., Ferguson, P. L., Browning, B. D., Bryant, B. E., Tomko, R. L., Gray, K. M., Mewton, L. & Squeglia, L. M. 2024. Predictors of Substance Use Initiation by Early Adolescence. Am J Psychiatry, 181, 423–433.

Hammerslag, L. R. & Gulley, J. M. 2016. Sex differences in behavior and neural development and their role in adolescent vulnerability to substance use. Behav Brain Res, 298, 15–26.

Jacobus, J. & Tapert, S. F. 2013. Neurotoxic effects of alcohol in adolescence. Annu Rev Clin Psychol, 9, 703–21.

Jin, C., Jia, H., Lanka, P., Rangaprakash, D., Li, L., Liu, T., Hu, X. & Deshpande, G. 2017. Dynamic brain connectivity is a better predictor of PTSD than static connectivity. Hum Brain Mapp, 38, 4479–4496.

Jones, C. M., Clayton, H. B., Deputy, N. P., Roehler, D. R., Ko, J. Y., Esser, M. B., Brookmeyer, K. A. & Hertz, M. F. 2020. Prescription Opioid Misuse and Use of Alcohol and Other Substances Among High School Students - Youth Risk Behavior Survey, United States, 2019. MMWR Suppl, 69, 38–46.

Jones, J. S., Monaghan, A., Leyland-Craggs, A., Team, C. & Astle, D. E. 2023. Testing the triple network model of psychopathology in a transdiagnostic neurodevelopmental cohort. Neuroimage Clin, 40, 103539.

Kaag, A. M., Wiers, R. W., De Vries, T. J., Pattij, T. & Goudriaan, A. E. 2019. Striatal alcohol cue-reactivity is stronger in male than female problem drinkers. Eur J Neurosci, 50, 2264–2273.

Kao, C. H., Khambhati, A. N., Bassett, D. S., Nassar, M. R., Mcguire, J. T., Gold, J. I. & Kable, J. W. 2020. Functional brain network reconfiguration during learning in a dynamic environment. Nat Commun, 11, 1682.

Kirse, H. A., Bahrami, M., Lyday, R. G., Simpson, S. L., Peterson-Sockwell, H., Burdette, J. H. & Laurienti, P. J. 2023. Differences in Brain Network Topology Based on Alcohol Use History in Adolescents. Brain Sci, 13.

Kullback, S. & Leibler, R. A. 1951. On Information and Sufficiency. Annals of Mathematical Statistics, 22, 79–86.

Kuntsche, E. & Muller, S. 2012. Why do young people start drinking? Motives for first-time alcohol consumption and links to risky drinking in early adolescence. Eur Addict Res, 18, 34–9.

Lees, B., Garcia, A. M., Debenham, J., Kirkland, A. E., Bryant, B. E., Mewton, L. & Squeglia, L. M. 2021. Promising vulnerability markers of substance use and misuse: A review of human neurobehavioral studies. Neuropharmacology, 187, 108500.

Marek, S. & Dosenbach, N. U. F. 2018. The frontoparietal network: function, electrophysiology, and importance of individual precision mapping. Dialogues Clin Neurosci, 20, 133–140.

Menon, V. 2011. Large-scale brain networks and psychopathology: a unifying triple network model. Trends Cogn Sci, 15, 483–506.

Mokhtari, F., Rejeski, W. J., Zhu, Y., Wu, G., Simpson, S. L., Burdette, J. H. & Laurienti, P. J. 2018. Dynamic fMRI networks predict success in a behavioral weight loss program among older adults. Neuroimage, 173, 421–433.

Morris, T. P., Kucyi, A., Anteraper, S. A., Geddes, M. R., Nieto-Castanon, A., Burzynska, A., Gothe, N. P., Fanning, J., Salerno, E. A., Whitfield-Gabrieli, S., Hillman, C. H., Mcauley, E. & Kramer, A. F. 2022. Resting state functional connectivity provides mechanistic predictions of future changes in sedentary behavior. Sci Rep, 12, 940.

Muller-Oehring, E. M., Kwon, D., Nagel, B. J., Sullivan, E. V., Chu, W., Rohlfing, T., Prouty, D., Nichols, B. N., Poline, J. B., Tapert, S. F., Brown, S. A., Cummins, K., Brumback, T., Colrain, I. M., Baker, F. C., De Bellis, M. D., Voyvodic, J. T., Clark, D. B., Pfefferbaum, A. & Pohl, K. M. 2018. Influences of Age, Sex, and Moderate Alcohol Drinking on the Intrinsic Functional Architecture of Adolescent Brains. Cereb Cortex, 28, 1049–1063.

Newman, M. E. J. & Girvan, M. 2004. Finding and evaluating community structure in networks. Physical Review E, 69.

NIAAA. 2011. *Alcohol screening and brief intervention for youth: A practitioner’s guide* [Online]. [Accessed].

Norman, A. L., Pulido, C., Squeglia, L. M., Spadoni, A. D., Paulus, M. P. & Tapert, S. F. 2011. Neural activation during inhibition predicts initiation of substance use in adolescence. Drug Alcohol Depend, 119, 216–23.

O’connell, J. & Hojsgaard, S. 2011. Hidden Semi Markov Models for Multiple Observation Sequences: The mhsmm Package for R. Journal of Statistical Software, 39, 1–22.

Ogawa, S., Lee, T. M., Kay, A. R. & Tank, D. W. 1990. Brain magnetic resonance imaging with contrast dependent on blood oxygenation. Proc Natl Acad Sci U S A, 87, 9868–72.

Patrick, M. E. & Schulenberg, J. E. 2013. Prevalence and Predictors of Adolescent Alcohol Use and Binge Drinking in the United States. Alcohol Research-Current Reviews, 35, 193–200.

Peltier, M. R., Verplaetse, T. L., Mineur, Y. S., Petrakis, I. L., Cosgrove, K. P., Picciotto, M. R. & Mckee, S. A. 2019. Sex differences in stress-related alcohol use. Neurobiol Stress, 10, 100149.

Petit, G., Kornreich, C., Verbanck, P. & Campanella, S. 2013. Gender differences in reactivity to alcohol cues in binge drinkers: a preliminary assessment of event-related potentials. Psychiatry Res, 209, 494–503.

Pruim, R. H. R., Mennes, M., Buitelaar, J. K. & Beckmann, C. F. 2015a. Evaluation of ICA-AROMA and alternative strategies for motion artifact removal in resting state fMRI. Neuroimage, 112, 278–287.

Pruim, R. H. R., Mennes, M., Van Rooij, D., Llera, A., Buitelaar, J. K. & Beckmann, C. F. 2015b. ICA-AROMA: A robust ICA-based strategy for removing motion artifacts from fMRI data. Neuroimage, 112, 267–277.

Raichle, M. E. 2015. The brain’s default mode network. Annu Rev Neurosci, 38, 433–47.

Rubinov, M. & Sporns, O. 2010. Complex network measures of brain connectivity: uses and interpretations. Neuroimage, 52, 1059–69.

Schulte, M. T., Ramo, D. & Brown, S. A. 2009. Gender differences in factors influencing alcohol use and drinking progression among adolescents. Clin Psychol Rev, 29, 535–47.

Seo, D., Jia, Z. R., Lacadie, C. M., Tsou, K. A., Bergquist, K. & Sinha, R. 2011. Sex Differences in Neural Responses to Stress and Alcohol Context Cues. Human Brain Mapping, 32, 1998–2013.

Shappell, H., Caffo, B. S., Pekar, J. J. & Lindquist, M. A. 2019. Improved state change estimation in dynamic functional connectivity using hidden semi-Markov models. Neuroimage, 191, 243–257.

Shappell, H. M., Duffy, K. A., Rosch, K. S., Pekar, J. J., Mostofsky, S. H., Lindquist, M. A. & Cohen, J. R. 2021. Children with attention-deficit/hyperactivity disorder spend more time in hyperconnected network states and less time in segregated network states as revealed by dynamic connectivity analysis. Neuroimage, 229, 117753.

Shen, X., Tokoglu, F., Papademetris, X. & Constable, R. T. 2013. Groupwise whole-brain parcellation from resting-state fMRI data for network node identification. Neuroimage, 82, 403–15.

Shine, J. M., Bissett, P. G., Bell, P. T., Koyejo, O., Balsters, J. H., Gorgolewski, K. J., Moodie, C. A. & Poldrack, R. A. 2016. The Dynamics of Functional Brain Networks: Integrated Network States during Cognitive Task Performance. Neuron, 92, 544–554.

Shine, J. M. & Poldrack, R. A. 2018. Principles of dynamic network reconfiguration across diverse brain states. Neuroimage, 180, 396–405.

Silveira, S., Shah, R., Nooner, K. B., Nagel, B. J., Tapert, S. F., De Bellis, M. D. & Mishra, J. 2020. Impact of Childhood Trauma on Executive Function in Adolescence-Mediating Functional Brain Networks and Prediction of High-Risk Drinking. Biol Psychiatry Cogn Neurosci Neuroimaging, 5, 499–509.

Somerville, L. H. & Casey, B. J. 2010. Developmental neurobiology of cognitive control and motivational systems. Curr Opin Neurobiol, 20, 236–41.

Spear, L. P. 2013. Adolescent neurodevelopment. J Adolesc Health, 52, S7–13.

Squeglia, L. M., Ball, T. M., Jacobus, J., Brumback, T., Mckenna, B. S., Nguyen-Louie, T. T., Sorg, S. F., Paulus, M. P. & Tapert, S. F. 2017. Neural Predictors of Initiating Alcohol Use During Adolescence. Am J Psychiatry, 174, 172–185.

Sridharan, D., Levitin, D. J. & Menon, V. 2008. A critical role for the right fronto-insular cortex in switching between central-executive and default-mode networks. Proc Natl Acad Sci U S A, 105, 12569–74.

Stautz, K. & Cooper, A. 2013. Impulsivity-related personality traits and adolescent alcohol use: a meta-analytic review. Clin Psychol Rev, 33, 574–92.

Stevens, W. D. & Spreng, R. N. 2014. Resting-state functional connectivity MRI reveals active processes central to cognition. Wiley Interdiscip Rev Cogn Sci, 5, 233–45.

Studer, B., Pedroni, A. & Rieskamp, J. 2013. Predicting risk-taking behavior from prefrontal resting-state activity and personality. PLoS One, 8, e76861.

Vaghi, M. M., Moutoussis, M., Vasa, F., Kievit, R. A., Hauser, T. U., Vertes, P. E., Shahar, N., Romero-Garcia, R., Kitzbichler, M. G., Bullmore, E. T., Consortium, N. & Dolan, R. J. 2020. Compulsivity is linked to reduced adolescent development of goal-directed control and frontostriatal functional connectivity. Proc Natl Acad Sci U S A, 117, 25911–25922.

Vink, M., Zandbelt, B. B., Gladwin, T., Hillegers, M., Hoogendam, J. M., Van Den Wildenberg, W. P., Du Plessis, S. & Kahn, R. S. 2014. Frontostriatal activity and connectivity increase during proactive inhibition across adolescence and early adulthood. Hum Brain Mapp, 35, 4415–27.

Zhao, Q., Sullivan, E. V., Muller-Oehring, E. M., Honnorat, N., Adeli, E., Podhajsky, S., Baker, F. C., Colrain, I. M., Prouty, D., Tapert, S. F., Brown, S. A., Meloy, M. J., Brumback, T., Nagel, B. J., Morales, A. M., Clark, D. B., Luna, B., De Bellis, M. D., Voyvodic, J. T., Nooner, K. B., Pfefferbaum, A. & Pohl, K. M. 2021. Adolescent alcohol use disrupts functional neurodevelopment in sensation seeking girls. Addict Biol, 26, e12914.

