## Supplementary Data for "Triple network dynamics and future alcohol consumption in adolescents"

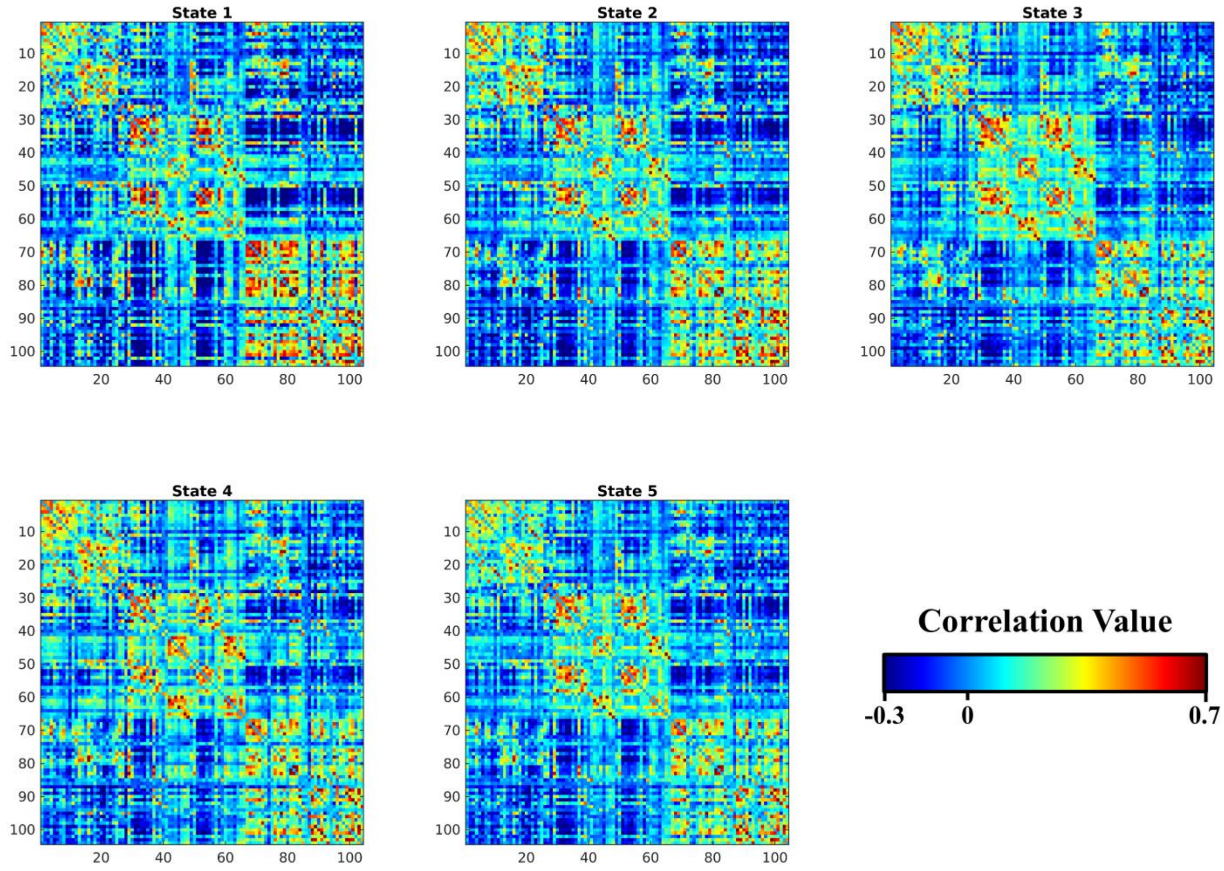


**Supplementary Figure 1. State covariance matrices**
Correlation matrices identified for each state by the Hidden Semi Markov Model. Color scales are the same across all matrices.

**Supplementary Table 1. NL versus RD sojourn distribution permutation testing**

| Brain State | $p_{unadjusted}$ |
|---|---|
| 1 | 0.666 |
| 2 | 0.621 |
| 3 | 0.591 |
| 4 | 0.486 |
| 5 | 0.881 |

Results are from permutation testing for differences in sojourn distribution between continual no/low (NL) versus future risky drinking (RD) participants. Each row shows results from a different brain state.

**Supplementary Table 2. Male versus female sojourn distribution permutation testing**

| Brain State | $p_{unadjusted}$ |
|---|---|
| 1 | 0.161 |
| 2 | **0.030** |
| 3 | 0.196 |
| 4 | 0.298 |
| 5 | 0.758 |

Results are from permutation testing for differences in sojourn distribution between continual male versus female participants. Each row shows results from a different brain state.

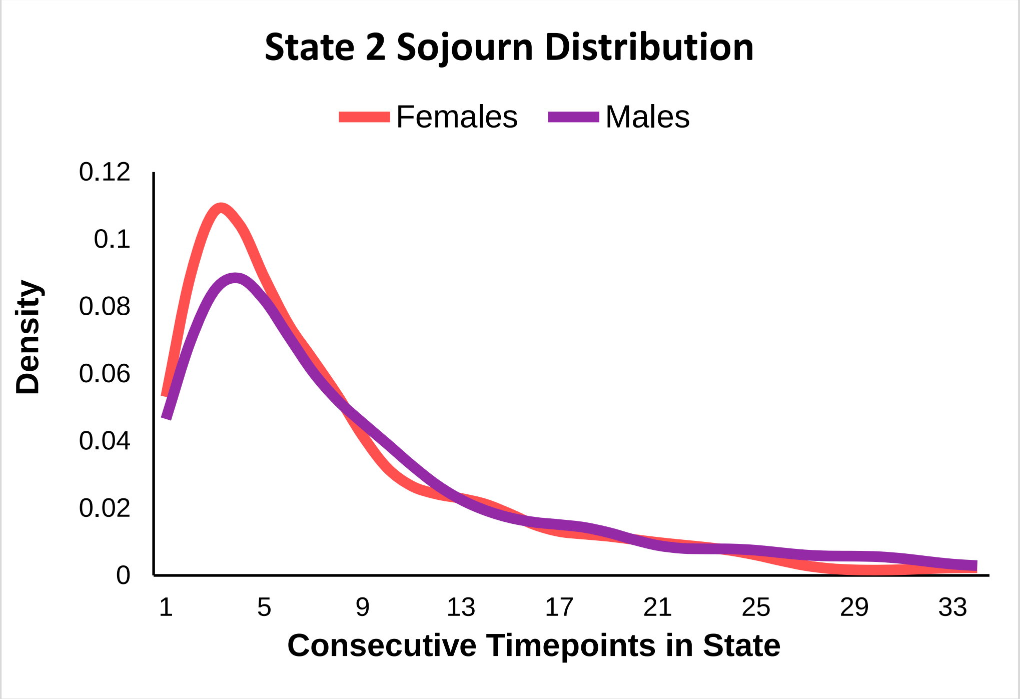


**Supplementary Figure 2. State 2 sojourn distributions – Male versus Female**
Density refers to the likelihood that a given sojourn into State 2 will last the number of consecutive TRs shown along the x-axis. After entering State 2, males tended to remain in the state for longer than females, though this effect was not statistically significant following multiple comparison correction for testing each of 5 states.

**Supplementary Table 3. Binomial Regression Results – Sex Interactions**

| Sequence Metric | β | Standard Error | $p_{Bonferroni}$ |
|---|---|---|---|
| Occupancy – State 1 | -0.0053 | 0.0057 | 1 |
| Occupancy – State 2 | 0.0040 | 0.0052 | 1 |
| Occupancy – State 3 | -0.0042 | 0.0047 | 1 |
| Occupancy – State 4 | 0.0043 | 0.0046 | 1 |
| Occupancy – State 5 | -0.0012 | 0.0066 | 1 |
| Number of Sojourns – State 1 | 0.0034 | 0.1132 | 1 |
| Number of Sojourns – State 2 | 0.1269 | 0.0791 | 1 |
| Number of Sojourns – State 3 | -0.0487 | 0.0913 | 1 |
| Number of Sojourns – State 4 | 0.0570 | 0.1015 | 1 |
| Number of Sojourns – State 5 | 0.0646 | 0.0723 | 1 |

Each row shows results from a different binomial regression model, all with drinking between visits (0 or 1) as the outcome variable. All models controlled for baseline age, time between visits, sex, socioeconomic status, study site, motion in fMRI scanner. Models with number of sojourns into states as the outcome of interest also controlled for the total number of sojourns a participant had throughout their fMRI scan. β-values indicate logistic relationships between independent and dependent variables.

**Supplementary Table 4. Binomial Regression Results – Main Effects**

| Sequence Metric | β | Standard Error | $p_{Bonferroni}$ |
|---|---|---|---|
| Occupancy – State 1 | 0.0015 | 0.0028 | 1 |
| Occupancy – State 2 | 0.0017 | 0.0027 | 1 |
| Occupancy – State 3 | 0.0002 | 0.0023 | 1 |
| Occupancy – State 4 | -0.0020 | 0.0023 | 1 |
| Occupancy – State 5 | -0.0028 | 0.0034 | 1 |
| Number of Sojourns – State 1 | 0.0100 | 0.0601 | 1 |
| Number of Sojourns – State 2 | 0.0039 | 0.0610 | 1 |

| | | | |
|---|---|---|---|
| Number of Sojourns – State 3 | 0.0167 | 0.0497 | 1 |
| Number of Sojourns – State 4 | -0.0366 | 0.0543 | 1 |
| Number of Sojourns – State 5 | 0.0050 | 0.0719 | 1 |

Each row shows results from a different binomial regression model, all with drinking between visits (0 or 1) as the outcome variable. All models controlled for baseline age, time between visits, sex, socioeconomic status, study site, motion in fMRI scanner. Models with number of sojourns into states as the outcome of interest also controlled for the total number of sojourns a participant had throughout their fMRI scan. β-values indicate logistic relationships between independent and dependent variables.
